# PrimPol primase mediates replication traverse of DNA interstrand crosslinks

**DOI:** 10.1101/2020.05.19.104729

**Authors:** Daniel González-Acosta, Elena Blanco-Romero, Karun Mutreja, Susana Llanos, Samuel Míguez, Fernando García, Javier Muñoz, Luis Blanco, Massimo Lopes, Juan Méndez

## Abstract

Interstrand crosslinks (ICLs) are DNA lesions frequently induced by chemotherapy that interfere with essential processes such as replication and transcription. ICL repair may be initiated by the convergence of two replication forks at the crosslink, which results in a termination-like DNA structure recognized and processed by the Fanconi Anemia (FA) pathway. An alternative possibility to generate a suitable substrate for ICL repair involves “ICL traverse”, a DNA damage tolerance mechanism in which a single fork arriving at the ICL can skip the lesion and restart DNA synthesis from a downstream point. This reaction requires FANCM translocase, the BLM/TOP3A/RMI1-2 (BTR) complex and other factors. Here we report that PrimPol, the second primase-polymerase identified in mammalian cells after Polα/Primase, interacts with BTR and participates in the ICL traverse reaction. A functional complementation assay reveals that the primase activity of PrimPol is required, confirming the need for re-priming events during ICL traverse. Genetic ablation of PRIMPOL strongly impaired this tolerance mechanism, making cells more dependent on fork convergence to initiate ICL repair. PRIMPOL KO cells and mice display hypersensitivity to ICL-inducing drugs, opening the possibility of targeting PrimPol activity to enhance the efficacy of chemotherapy based on DNA crosslinking agents.

## INTRODUCTION

DNA interstrand crosslinks (ICLs) are repaired in a DNA replication-dependent manner by a protein network that, if lost or altered by mutation, results in Fanconi Anemia (FA), a rare but very severe disease (reviewed by Ceccaldi et al, 2016). ICLs are recognized by the FANCM complex (FANCM/MHF1-2/FAAP24), which assists in the recruitment of the FA core complex responsible for the ubiquitylation and activation of the FANCD2/I heterodimer (Deans and West, 2009). Activated FANCD2/I recruits specific nucleases to one of the DNA strands initiating a complex process that involves homologous recombination (HR), translesion synthesis (TLS) and nucleotide excision repair (NER) mechanisms (reviewed by Deans and West, 2011; Zhang and Walter, 2014; Lopez-Martinez et al., 2016).

Many mechanistic aspects of the initial steps in ICL repair have been derived from biochemical studies in *X. laevis* egg extracts (Räschle et al., 2008; Knipscheer et al., 2009). In this system, ICL repair is triggered by the convergence of two forks that reach the ICL from opposite sides, creating an X-shaped structure that resembles a replication termination event (Zhang et al., 2015a). In mammalian cells, an alternative mechanism has been proposed that involves the bypass of the ICL by the first fork reaching the lesion. This reaction has been called ICL ‘traverse’ and is a form of DNA damage tolerance (DDT; Huang et al., 2013). Traverse reactions may account for up to 60% of the replication events monitored around ICLs in human cells and require FANCM (Huang et al., 2013), its interaction with PCNA (Rohleder et al., 2016) and the BTR complex (BLM/TOP3A/RMI1-2; Ling et al., 2016). The loss of BLM helicase activity within the BTR reduced the frequency of ICL traverse in chicken DT40 cells, suggesting that BLM might unwind the DNA at the unreplicated side of the ICL (Ling et al., 2016). The accumulation of ICLs also triggers an ATR-mediated global slowdown of replication forks and fork remodeling events that assist the traverse reaction (Mutreja et al., 2018).

The molecular events that take place during the traverse of ICLs are starting to be elucidated. It has been recently reported that ATR signaling and FANCD2 protein mediate the interaction between FANCM and the CMG helicase (Cdc45-MCM-GINS) and trigger the eviction of its GINS component. This remodeling of CMG may facilitate the opening of the MCM hexameric ring, a necessary step for the translocation of the replisome across the ICL (Huang et al., 2019). After the replisome is relocated downstream of the ICL, restart of DNA synthesis by DNA polymerases strictly requires re-priming events that to date have not been characterized. Besides the canonical Polα/Primase involved in DNA replication, many archaeal and eukaryotic organisms encode PrimPol, a primase-polymerase specialized in DNA damage tolerance (Bianchi et al., 2013; García-Gómez et al., 2013; Mourón et al., 2013; Wan et al., 2013). Human PrimPol is distributed between cell nuclei, cytosol and mitochondria and it is rapidly concentrated to chromatin upon UVC irradiation, to re-prime DNA synthesis downstream of UVC-induced photoproducts. Re-priming reactions allow fork progression and leave bypassed lesions to be repaired post-replicatively (Mourón et al., 2013). A related function for PrimPol during mitochondrial DNA synthesis was proposed (García-Gómez et al., 2013) and has been recently supported by analysis of replication intermediates in response to several stressors (Torregrosa-Muñumer et al., 2017). The interaction of PrimPol with ssDNA-binding proteins RPA and mitochondrial SSBP1 facilitates its recruitment to ssDNA generated next to DNA lesions (Wan et al., 2013; Guilliam et al., 2015; 2017). Furthermore, long ssDNA stretches enhance the primase activity of PrimPol *in vitro* (Martínez-Jiménez et al., 2017). The replicative tolerance function of PrimPol also operates at natural replication blocks formed by special structures such as G-quadruplexes (Schiavone et al, 2016) and R-loops (Švikovic et al, 2019).

Here we show that human cells rely on PrimPol primase activity to mediate the ICL traverse reaction, sustaining DNA synthesis during the cellular recovery from ICL-inducing agents. In the absence of PRIMPOL, cells become more dependent on the fork convergence mechanism, and the global efficiency of ICLs repair is compromised.

## RESULTS

### PrimPol interacts with ICL recognition and repair factors

A search for new PrimPol-interacting factors was conducted by immunoprecipitation (IP) coupled to mass spectrometry in U2OS cells synchronized in S phase. As control for non-specific binding proteins, we used U2OS PRIMPOL KO cells generated by CRISPR/Cas9 technology (**Figure 1A and Supp. Figure 1**). A quantitative proteomic analysis (enrichment ratio and statistical significance of proteins identified in WT vs KO cells) confirmed the efficient IP of PrimPol and the co-precipitation of known interacting factors RPA and SSBP1 (**Figure 1B**). Amongst the new potential PrimPol-interacting proteins were HERC2, involved in the resolution of G4 quadruplex structures (Wu et al., 2018) and several proteins linked to the recognition and repair of ICL lesions: MHF1 (part of the FANCM complex; Ciccia et al., 2007; Yan et al., 2010; Wang et al., 2013); RUNX1 (Tay et al., 2018); and BLM, RMI1 and RMI2 (three components of the BTR complex; Manthei and Keck, 2013). Both MHF1 and BTR participate in the ICL traverse reaction (Huang et al., 2013; Ling et al., 2016). The interaction between PrimPol and BLM, RMI1 and RMI2 was confirmed using IP-immunoblot assays, which also revealed the co-precipitation of TOP3A, the fourth component of the BTR complex (**Figure 1C**).

**Figure 1.**
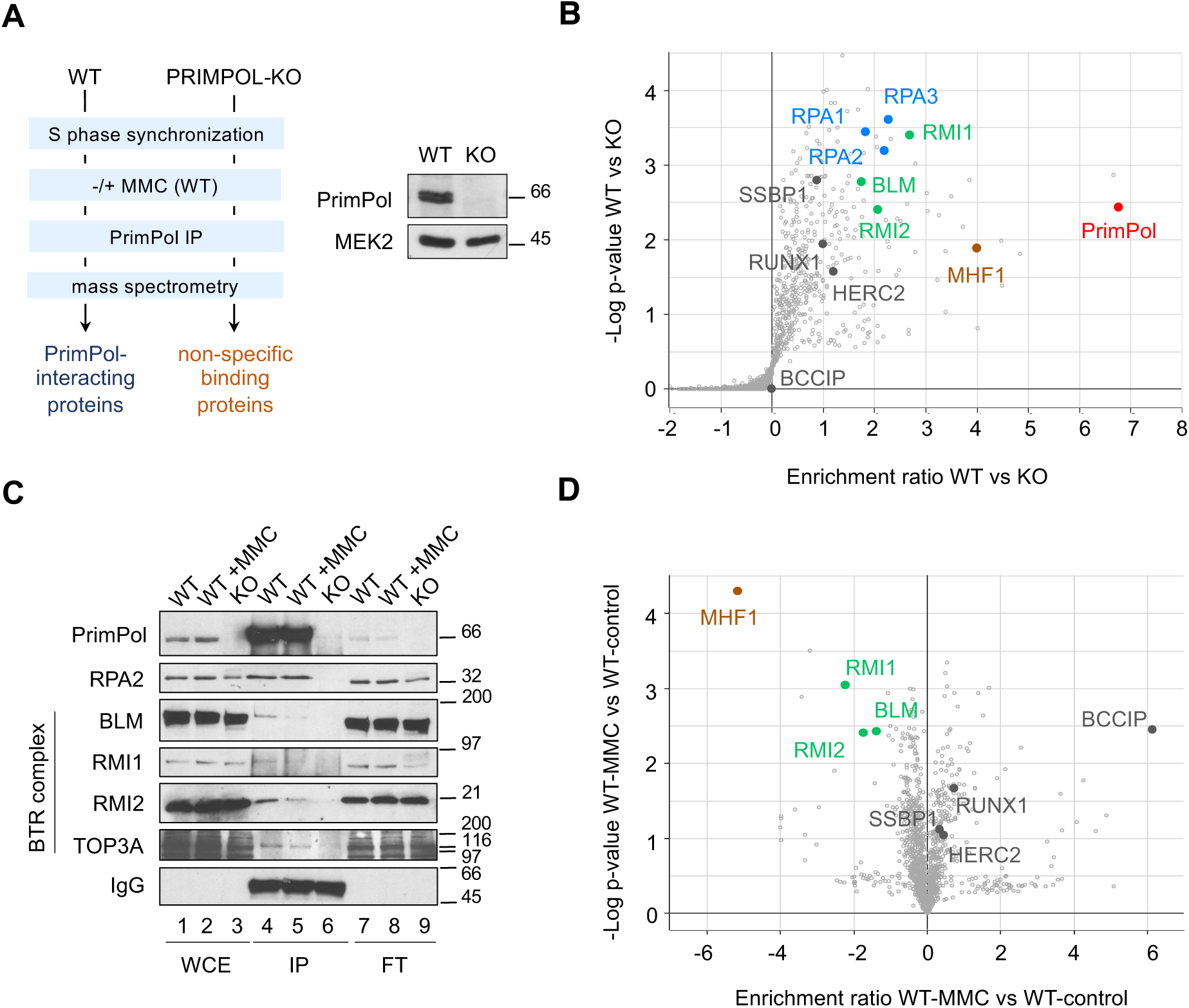
PrimPol interacts with factors involved in ICL traverse. **A**. Experimental design for the proteomic identification of PrimPol-interacting factors in S phase-synchronized cells. Immunoblots showing PrimPol levels in WT and KO cells. MEK2 is shown as loading control. **B**. Volcano plot showing statistical significance (-log p-value) vs enrichment ratio (log2) of identified proteins after PrimPol immunoprecipitation in WT and KO cells. The positions of PrimPol, RPA1-3, BLM, RMI1, RMI2, MHF1, SSBP1, RUNX1, HERC2 and BCCIP proteins are indicated. **C**. Immunoblot detection of the indicated proteins in whole cell extracts (WCE), PrimPol immunoprecipitates (IP), and non-precipitated flow-through fractions (FT). IgG shows the presence of anti-PrimPol antibody in the IP fractions (lanes 4-6). **D**. Volcano plot showing statistical significance (-log p-value) vs enrichment ratio (log2) of identified proteins after PrimPol immunoprecipitation in WT cells with or without MMC. BLM, RMI1, RMI2, MHF1, SSBP1, RUNX1, HERC2 and BCCIP proteins are indicated.

The IP-mass spectrometry assay was also performed in cells treated with mitomycin C (MMC), a DNA crosslinking agent used in chemotherapy. The comparison of MMC-treated vs control cells hinted at dynamic changes in the association between PrimPol and some of its potential partners. Interactions with SSBP1, HERC2, RUNX and the homologous recombination protein BCCIP (Lu et al., 2005) were enhanced, whereas the binding to BLM, RMI1, RMI2 and MHF1 was apparently reduced (**Figure 1D, Supp. Figure 1C**). The effect of MMC on the PrimPol-BTR interaction was validated by IP-immunoblots (**Figure 1C**, lanes 4 and 5) and suggests that the physical interaction between PrimPol and the BTR complex is disrupted during the cellular response to MMC.

### PrimPol facilitates DNA synthesis in response to ICL-inducing agents

Following UV-C irradiation or treatment with cisplatin, PrimPol accumulates on chromatin to facilitate the replicative bypass of DNA photoadducts or cisplatin-induced lesions (Mourón et al., 2013; Quinet et al., 2020). To test whether PrimPol is recruited to chromatin in response to ICLs, we analyzed its subcellular localization in response to MMC (that induces DNA monoadducts, intra-strand crosslinks and ICLs), or trimethyl-psoralen activated with UVA (TMP-UVA), which induces ICLs with high specificity (∼90% of total lesions; Lopez-Martinez et al., 2016). Both treatments induced the accumulation of PrimPol protein on chromatin (**Figure 2A**, lanes 9-12). Ubiquitylation of FANCD2 protein and its enrichment on chromatin indicated the activation of the ICL repair pathway in both cases. This result suggests that PrimPol participates in the replicative tolerance and/or repair of ICLs.

**Figure 2.**
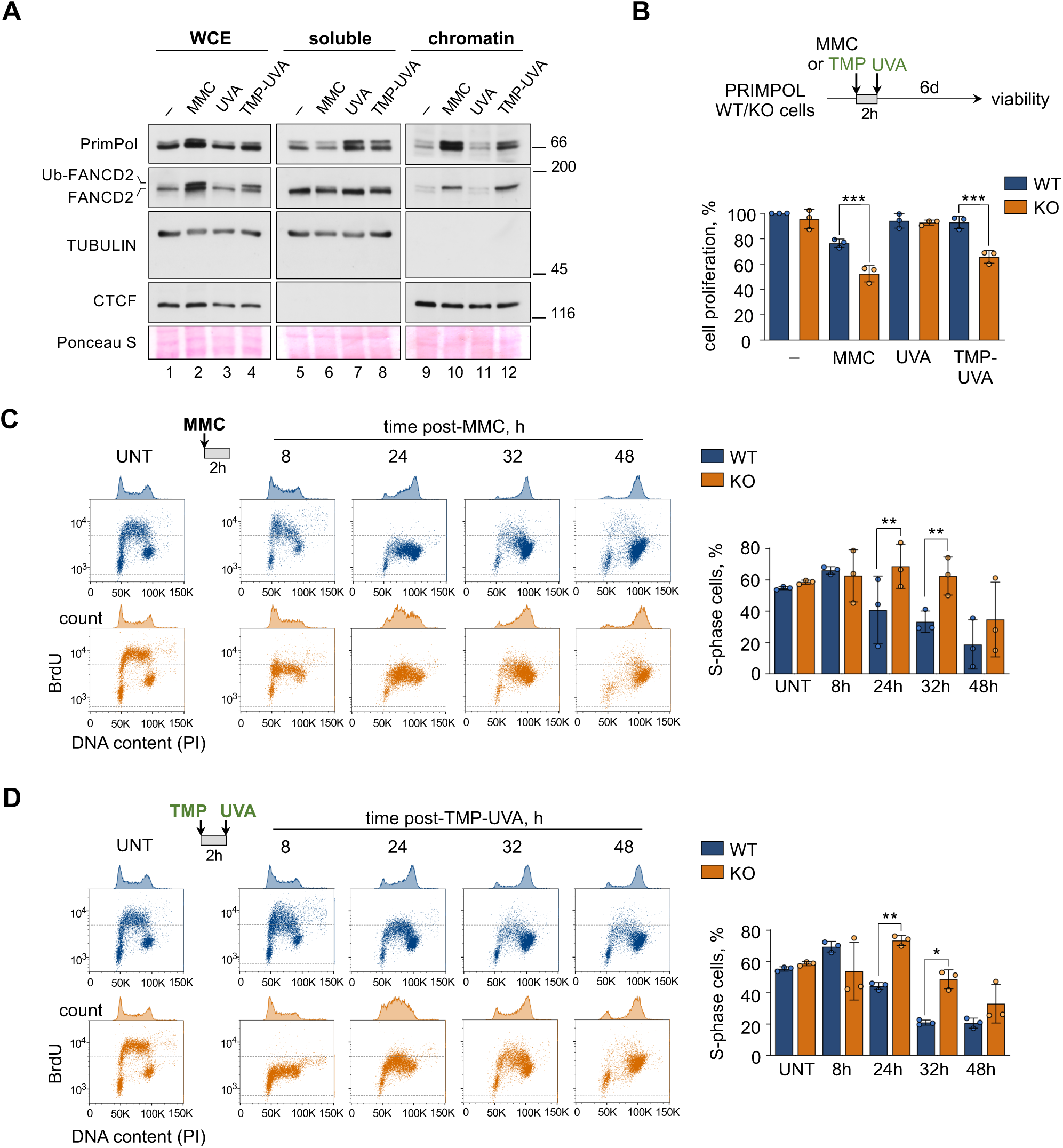
PrimPol facilitates DNA synthesis and cell proliferation in the presence of ICLs. **A**. Immunoblots show the levels of the indicated proteins in whole cell extracts (WCE), soluble and chromatin-enriched fractions. TUBULIN and CTCF are shown as controls for soluble and chromatin-bound proteins, respectively. The position of the ubiquitylated form of FANCD2 (Ub-FANCD2) is indicated. Ponceau-S staining is shown as a loading control. **B**. Cell viability assays in WT and PRIMPOL KO cells treated with 1 µg/ml MMC (2 h) or TMP-UVA (2 µM TMP, 2h followed by 5 s irradiation), and incubated for 6 d in regular medium. Viability was assessed with CellTiter-Glo. Histogram represents the average and SD of three assays. Circle dots in each column represent the values of individual replicates. Statistical analysis was conducted with one-way ANOVA and Bonferroni post-test. ***, p<0.001. **C**. Flow cytometry profiles of BrdU incorporation (y-axis) vs DNA content (PI; x-axis) in WT (blue) and PRIMPOL KO (orange) cells, either collected without treatment (UNT) or at the indicated times after exposure to 1 µg/ml MMC (2 h). DNA content profiles are shown on top of the boxes (cell count). Dashed horizontal lines inside panels are included for comparative purposes between WT and KO BrdU incorporation levels. Right, histograms represent the percentage of cells in S-phase in each condition (average and SD of three assays). Circle dots in each column represent the values of individual replicates. **D**. Same as (C) except that the treatment was TMP-UVA (2 µM TMP for 2h followed by 5 s irradiation). Statistical analysis in all cases was conducted with one-way ANOVA and Bonferroni post-test. *, p<0.05; **, p<0.01.

To assess the relevance of PrimPol in the cellular response to ICLs, WT and PRIMPOL KO cells were treated with MMC or TMP for 2h (TMP was activated with UVA right after incubation), and kept in regular media for 6 days to estimate their proliferation capacity and viability. The number of living cells at the end of the experiment was significantly reduced in PRIMPOL KO cells, reflecting a higher sensitivity to the DNA damage inflicted by either treatment. The drop in viability was not observed with UVA in the absence of TMP, suggesting that the effect was mediated by ICL generation (**Figure 2B**). A similar hypersensitivity to MMC or TMP-UVA occurred when PRIMPOL was downregulated with shRNA, arguing against the possibility of off-target effects caused by the CRISPR/Cas9 genetic ablation (**Supp. Figure 2A**).

We next tested whether the increased sensitivity of PrimPol KO cells was related to the dynamics of DNA replication following the exposure to crosslinking agents. Untreated WT and PRIMPOL KO cells displayed virtually identical DNA content and BrdU incorporation profiles (**Figure 2C-D**, left panels). Upon administration of MMC or TMP-UVA for 2h to an asynchronous cell population, DNA content analyses revealed a slow progression through the cell cycle, requiring up to 32 h for full completion of S phase (**Figure 2C-D**, blue profiles). In PrimPol KO cells, this effect was exacerbated and S phase completion required up to 48 h (**Figure 2C-D;** orange profiles). As expected, UVA did not affect cell cycle progression of WT or KO cells in the absence of TMP (**Supp. Figure 2B**). The effects of MMC and TMP-UVA in WT and KO cells were confirmed by flow cytometry profiles of BrdU incorporation (**Figure 2C-D**) and by confocal microscopy analysis of EdU incorporation (**Supp. Figure 2C-D**). Of note, this phenotype was rescued by the reintroduction of exogenous PrimPol in KO cells (**Supp. Figure 2C-E**). We conclude from these experiments that PrimPol promotes continuous DNA synthesis during the initial response to DNA damage induced by MMC and TMP-UVA.

### PrimPol restarts DNA synthesis at forks stalled by crosslinking agents

The slower progression through S phase in PRIMPOL KO cells following MMC or TMP-UVA treatments might derive from an inefficient restart of DNA synthesis in forks stalled at DNA lesions. To test this hypothesis, we estimated the frequency of stalled and restarted forks in response to MMC. WT and PRIMPOL KO cells were pulse-labeled with CldU (20’), treated with MMC (30’) and then pulse-labeled with IdU (20’). Upon DNA fiber stretching and immunolabeling with anti-CldU (red) and anti-IdU (green) antibodies, forks stalled at MMC-induced lesions can be divided in those that remained inactive for the duration of the experiment (red tracks), and those that restarted DNA synthesis (red-green tracks). Non-stalled forks were also identified by the presence of a gap between the red and green signals (Mourón et al., 2013; **Figure 3A**). Using this assay, we observed that the percentage of fork restart in WT cells was severely reduced in PRIMPOL KO cells (80% vs 40%; **Figure 3B**). This phenotype was rescued by the reintroduction of V5-tagged PrimPol in KO cells, but not by a catalytic mutant defective in both primase and polymerase activities (AxA; García-Gómez et al., 2013). A PrimPol mutant (ΔZn) that is primase-deficient but polymerase-proficient (García-Gómez et al., 2013; Mourón et al., 2013) also failed to rescue the defect in fork restart (**Figure 3B-C and Supp. Figure 3**), indicating that the primase activity of PrimPol mediates the re-initiation of DNA synthesis at forks stalled by MMC.

**Figure 3.**
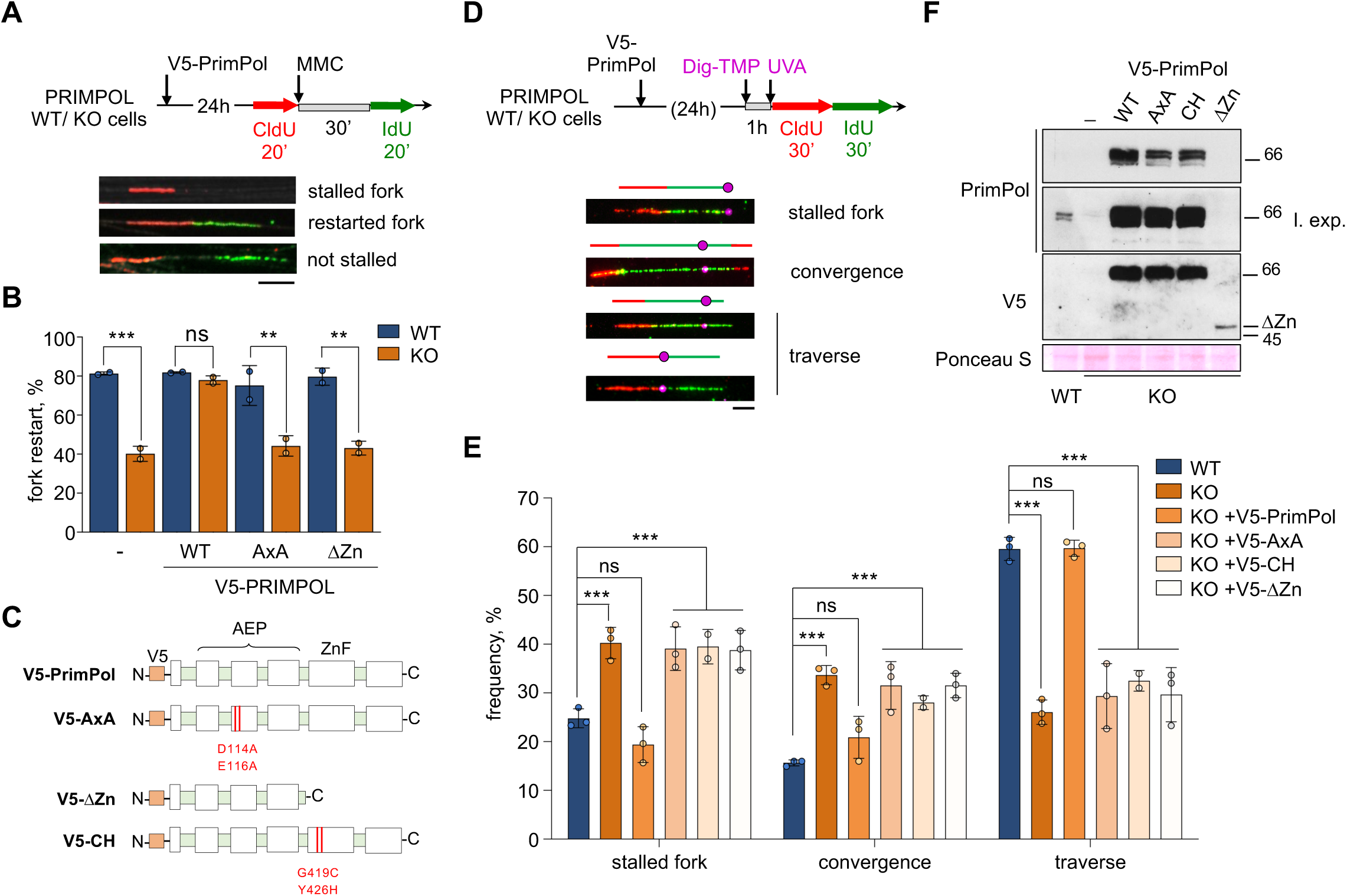
PrimPol mediates the ICL traverse reaction through its primase activity. **A**. Schematic of the experimental design. Examples of stalled, restarted forks and a fork not stalled by MMC lesion. Scale bar, 10 µm. **B**. Histogram shows the percentage of fork restart (average and SD of two replicates) in each condition. Circle dots in each column represent the values of individual replicates. Statistical analysis was conducted with one-way ANOVA and Bonferroni post-test. **, p<0.01; ***, p<0.001; ns, not significant. WT indicates the expression of V5-PrimPol protein; AxA, a PrimPol catalytic mutant; ΔZn, a primase-null, polymerase-proficient PrimPol mutant. **C**. PrimPol mutants used in the experiments. **D**. Experimental design and examples of stretched DNA fibers with different patterns of DNA synthesis around the ICL lesion. Fibers were stained with anti-CldU (red), anti-IdU (green) and anti-Dig (magenta). Schematics under the fiber images highlight the position of the ICL (circle) in each replicative structure. **E**. Histograms show the percentage of the different replication patterns observed in each experimental condition (average and SD of three assays). Circle dots in each column represent the values of individual replicates. AxA, PrimPol catalytic mutant. ΔZn and CH are primase-null, polymerase-proficient PrimPol mutants. Statistical analysis: two-way ANOVA followed by Bonferroni post-test comparing each condition to WT. ***, p<0.001; ns, not significant. **F**. Immunoblots showing the levels of V5-PrimPol proteins (WT and mutant derivatives) used in (E). Ponceau-S staining is shown as loading control.

### PrimPol mediates ICL traverse reactions

Because not all crosslinks induced by MMC are ICLs, the effect in fork restart described above could reflect PrimPol-mediated bypass of intra-strand adducts or other types of lesions. To test directly whether PrimPol re-primes DNA synthesis downstream of ICLs, we used TMP molecules linked to digoxigenin (Dig-TMP) to localize the positions of ICLs in stretched DNA fibers by immunofluorescence with anti-Dig antibody (Huang et al., 2013; Mutreja et al., 2018). This approach allows the identification of three main types of replication tracks around ICLs: (1) single forks stalled at the ICL; (2) two forks converging at the ICL; (3) single forks that have traversed the lesion (**Figure 3D**). In WT cells, the majority of labeled tracks corresponded to ICL traverse reactions (60%), followed by single stalled forks (25%) and converged forks (15%), in agreement with previous reports (Huang et al., 2013; 2019; **Figure 3E**). In PRIMPOL KO cells, however, the frequency of traverse reactions was drastically reduced (26%, less than half than WT cells), while fork convergence events increased to 34%. This effect could be entirely attributed to PrimPol protein, as reintroduction of exogenous PrimPol in KO cells restored the proportions observed in WT cells. Neither the catalytic mutant (AxA) nor the primase-deficient ΔZn version of PrimPol could rescue the frequency of ICL traverse reactions (**Figure 3E-F**), underscoring the importance of the primase activity. Because the ΔZn mutant was expressed at lower levels than WT protein (**Figure 3F** and **Supp. Figure 3B**), we confirmed this result using PrimPol-CH, a full-length protein carrying two mutations in the Zn-finger domain (C419G/H426Y) that inactivate the primase (Mourón et al., 2013; **Figure 3C, E-F**). These assays provide molecular evidence that PrimPol mediates the ICL traverse reaction, and this function strictly requires its primase activity.

### Defective ICL repair and chromosomal instability in PRIMPOL KO cells

Next, we tested whether the lower frequency of ICL traverse reactions observed in PRIMPOL KO cells affected the dynamics of nuclear foci of FANCD2 protein, a marker of active ICL repair (Garcia-Higuera et al., 2001; Liang et al., 2015). In WT cells, FANCD2 foci were apparent as early as 8 h post-TMP-UVA (average 19 foci/cell), but were barely detected in KO cells. By 24 h, both WT and KO cells accumulated FANCD2 foci (av. 35-40 foci/cell). By 48 h, the number of foci was significantly reduced in WT cells (av. 15 foci/cell) but remained high in KO cells (av. 25 foci/cell; **Figure 4A-B**). The differences in FANCD2 foci dynamics between WT and KO cells were also observed after exposure to MMC (**Supp. Figure 4**). Of note, karyotypic analyses in metaphase spreads revealed the accumulation of chromosomal breaks, gaps and fusions preferentially in PRIMPOL KO cells treated with TMP-UVA (**Figure 4C-D**), which could result from cells having entered mitosis with unrepaired ICLs (Akkari et al., 2000; Mutreja et al., 2018). Combined, these results indicate a delayed initiation and apparently incomplete ICL repair process in the absence of PrimPol protein.

**Figure 4.**
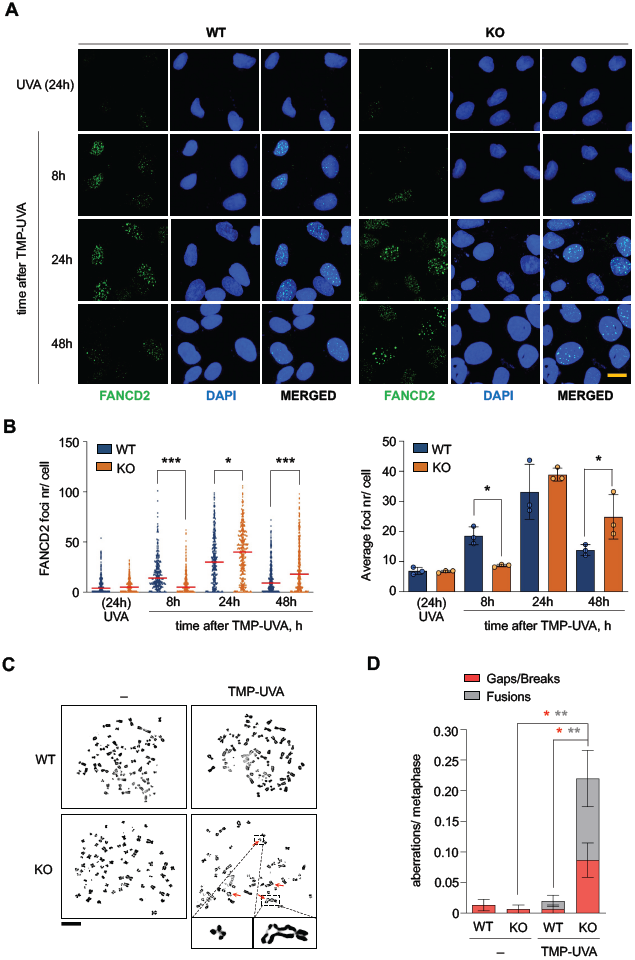
ICL repair is impaired upon loss of PrimPol. **A**. Confocal microscopy images of FANCD2 immunofluorescence staining in control (UVA) or TMP-UVA-treated (2 µM TMP, 2h followed by 5 s irradiation) WT and PRIMPOL KO cells. Nuclear DNA is counterstained with DAPI. Scale bar, 25µm. **B**. Dot plot shows the distribution and mean (horizontal red line) of FANCD2 foci number per cell in each condition. Data pooled from three replicates are represented (n ≥100 cells per condition). Statistical analysis was conducted with Kruskal-Wallis test and Dunns post-test. *, p<0.05; ***, p<0.001. Histograms (right panel) show the average foci number and SD of three assays in each case. Circle dots in each column represent the values of individual replicates. Statistical analysis was conducted with one-way ANOVA followed by Bonferroni post-test. *, p<0.05. **C**. Metaphase spreads stained with Leishman, obtained from control (-) or TMP-UVA-treated cells (2 µM TMP, 2h followed by 5 s irradiation). Scale bar, 10 µm. Representative images of chromosomal breaks and fusions are shown at higher magnification. **D**. Histograms show the number of chromosomal aberrations per metaphase (average and SEM) in each condition. Data taken from three replicates are combined (50 metaphases/ replicate). Statistical analysis was conducted with one-way ANOVA followed by Bonferroni post-test. *, p<0.05; **, p<0.01.

### Hypersensitivity of PrimPol KO mice to MMC

Given its roles in DNA damage tolerance, now including the traverse of ICLs, it could be predicted that PrimPol protects cells from the toxicity of DNA crosslinking agents. In this regard, PRIMPOL KO cells displayed hypersensitivity to cisplatin (Quinet et al., 2020) and MMC (**Figure 5A**). To test the relevance of this protective function *in vivo*, WT and PRIMPOL KO mice were treated with different concentrations of MMC. In mice, MMC induces a rapid depletion of haematopoietic progenitor cells in the bone marrow (BM) that is exacerbated in strains deficient for FAN1, MUS81, SNM1 and FAAP20, all required for ICL repair (Dendouga et al., 2005; Dronkert et al., 2000; McPherson et al., 2004; Thongthip et al., 2016; Zhang et al., 2015b). A dose of 7.5 mg MMC/kg of body weight was innocuous for WT mice but lethal for 20% of KO mice. At a higher MMC concentration (10 mg/kg), the median survival was 13 days in WT mice and 10 days in KO mice. At 15 mg/kg, the median survival of KO mice was 4 days shorter than WT (7 vs 11 days; **Figure 5B**).

**Figure 5.**
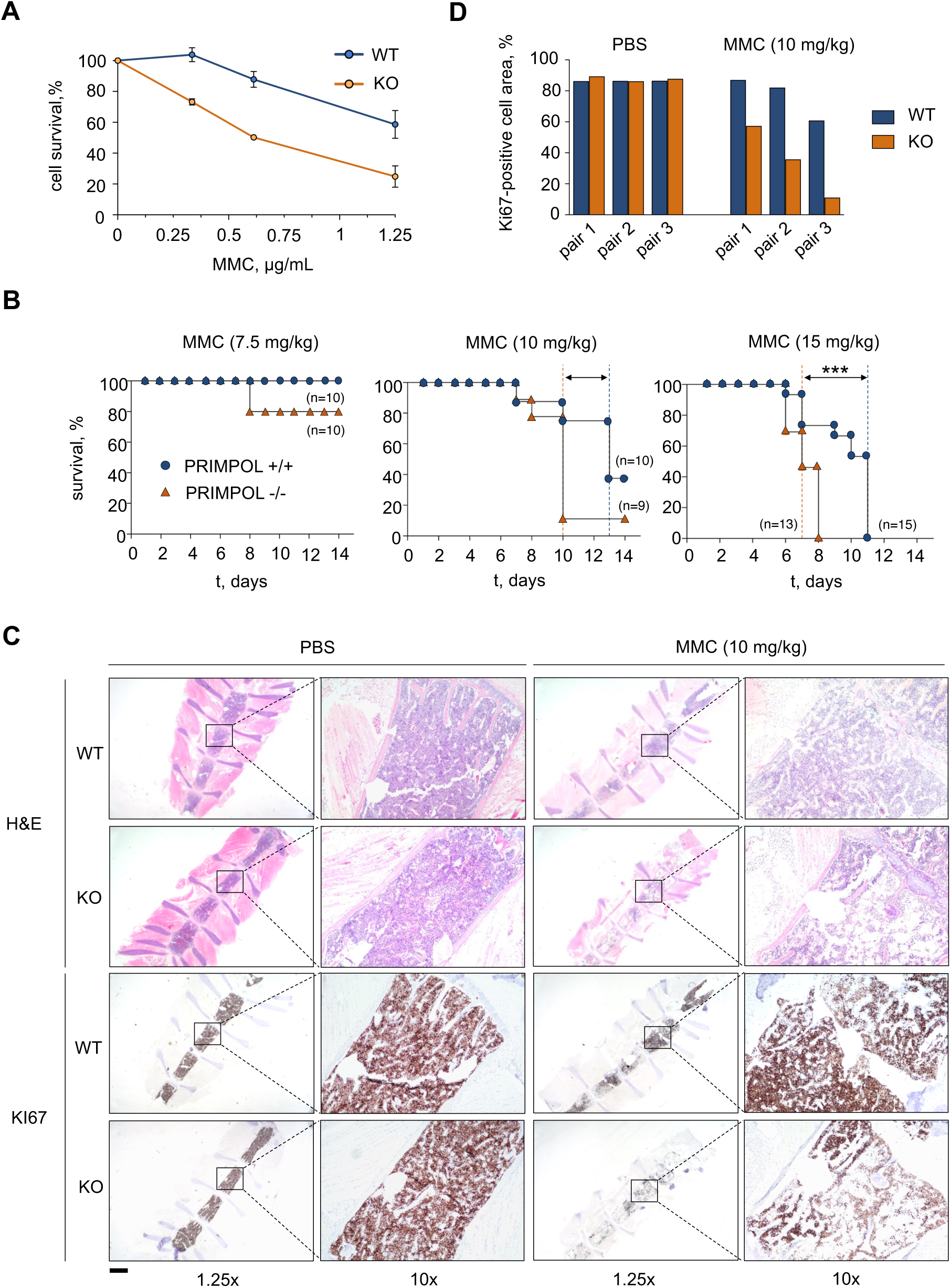
PRIMPOL KO cells and mice are hypersensitive to MMC. **A**. Survival curves of WT and PRIMPOL KO cells cultured for 72 h in the presence of increasing amounts of MMC. Cell survival was assessed with CellTiter-Glo. Each point in the curve represents the percentage of surviving cells (average and SD of three assays). **B**. Kaplan-Meier survival curves of WT and PRIMPOL KO mice after the administration of a single dose of MMC (7.5, 10 or 15 mg/kg). Dashed vertical lines represent median survival values for WT (blue) and KO (orange) mice. **C**. Haematoxylin-eosin (H&E) and Ki-67 immunohistochemistry (IHC) stainings of bone marrow sections derived from WT and PRIMPOL KO mice following the administration of 10 mg/kg MMC. Control mice (PBS) were sacrificed when MMC-injected mice reached the humane endpoint. Scale bar, 2.5 mm. In each image, a relevant area is shown at higher magnification (10x). **D**. Histograms show Ki-67-positive cellular area in the conditions shown in (C). Results are shown with three pairs of age- and gender-matched WT and KO littermates. When combined, the differences observed between MMC-treated WT and KO mice were statistically significant (two-way ANOVA and Bonferroni post-test, p-value 0.0044).

To compare the proliferation status of BM cells, three pairs of age- and gender-matched WT and KO mice were treated with 10 mg/kg MMC. As KO mice are more sensitive to MMC, WT individuals were euthanized when their paired KO mice had reached the humane endpoint, and their BMs were analyzed in parallel (**Figure 5C**). In untreated mice, the BM of WT and KO displayed similar cellularity and proliferation rate. In contrast, in response to MMC, the percentage of Ki67-positive cells was reduced in the BM of KO mice compared to WT (**Figure 5C-D**). This result confirms the protective function of PrimPol in response to DNA crosslinking agents *in vivo*.

### Activation of extra origins alleviates the loss of PrimPol

Because the ICL traverse reaction is impaired in PRIMPOL KO cells, they may become more dependent on the fork convergence mechanism to trigger ICL repair. The probability of a second fork reaching the ICL is increased by the activation of dormant origins, whose availability depends on the cellular concentration of MCM proteins (Ge et al., 2007; Ibarra et al., 2008). To identify possible synthetic effects between the loss of PrimPol and the defective activation of dormant origins, MCM3 expression was downregulated with RNAi (**Figure 6A)**. The reduction in the cellular levels of Mcm3 protein increased the sensitivity of PRIMPOL KO cells to MMC to a higher extent than WT cells (**Figure 6B**). To demonstrate that Mcm3 downregulation prevented the activation of backup origins, we estimated origin activity in DNA fibers by measuring the abundance of origin tracks relative to other replicative structures, e.g. forks and termination events (**Figure 6C**). Even in an unchallenged S phase, PrimPol KO cells displayed a higher frequency of origin activation, as previously reported (**Figure 6D**, columns 1 and 5; Rodriguez-Acebes et al., 2018). Treatment with MMC increased origin activity in both WT and KO cells, reflecting the activation of backup, dormant origins (**Figure 6D**, columns 1-2 and 5-6). As expected, MCM3 downregulation did not affect origin activation in unchallenged conditions (**Figure 6D**, compare 1 with 3 and 5 with 7), but it completely prevented the activation of additional origins in both WT and KO cells upon MMC treatment (**Figure 6D**, compare 3 with 4, and 7 with 8).

**Figure 6.**
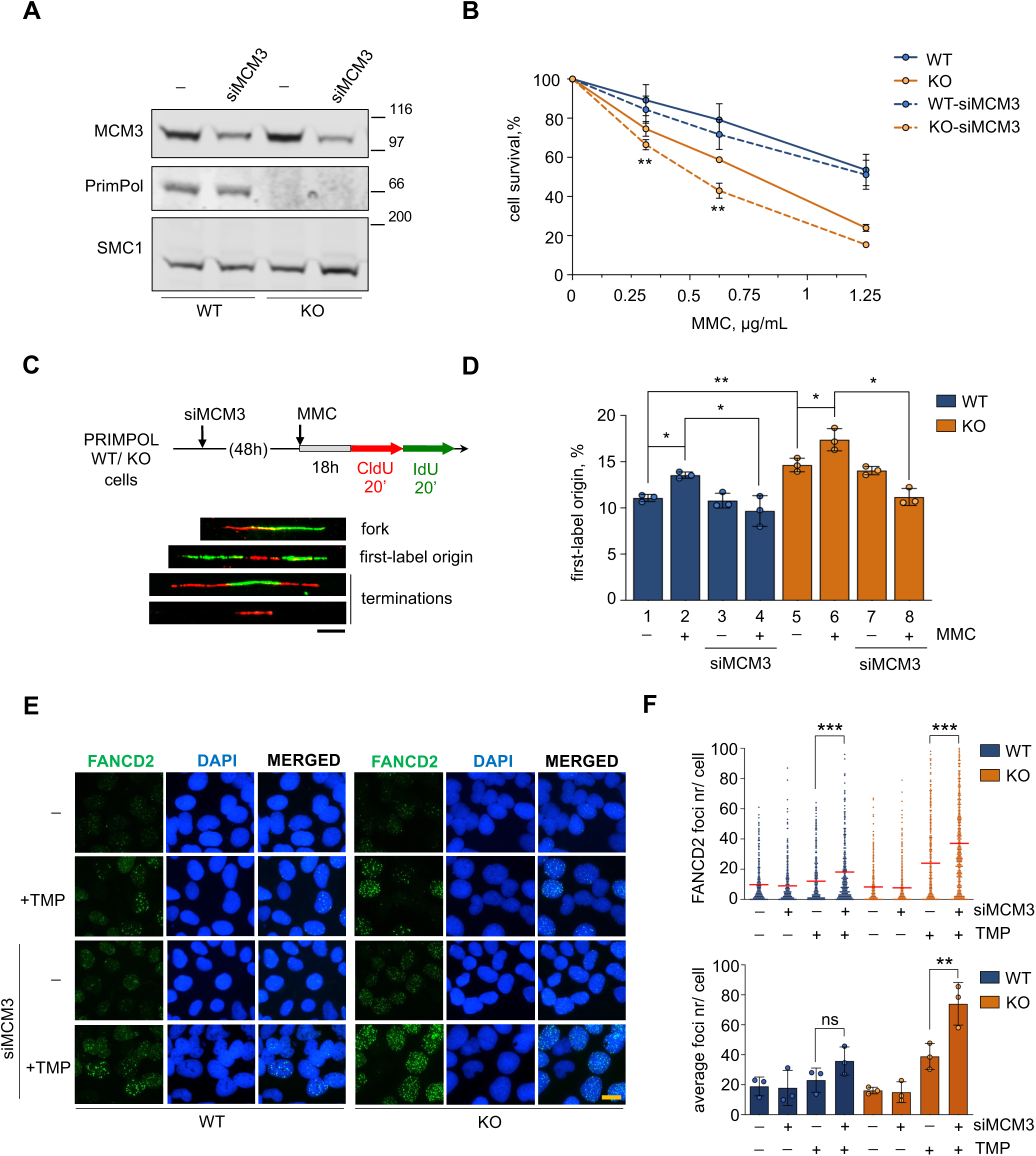
Loss of backup origins exacerbates the effect of PrimPol loss in ICL repair. **A**. Immunoblots showing the levels of MCM3 and PrimPol proteins in PRIMPOL WT and KO cells. SMC1 is shown as loading control. **B**. Cell survival assays with WT and PRIMPOL KO cells following MCM3 downregulation with siRNA in the presence of increasing amounts of MMC for 72 h. Each point in the curve represents the percentage of surviving cells (average and SD of three assays). Statistical analysis was conducted with two-way ANOVA followed by Bonferroni post-test. (**, p<0.01; comparison of KO and KO-siMCM3). **C**. Schematic of experimental design and examples of replicative structures (fork/ first-label origin/ terminations) in stretched DNA fibers. Scale bar, 10 µm. **D**. Percentage of first-label origins in each condition (average and SD of three assays). Circle dots in each column represent the values of individual replicates. Statistical analysis: one-way ANOVA followed by Bonferroni post-test. *, p<0.05; **, p<0.01. **E**. IF detection of FANCD2 foci in WT and PRIMPOL KO cells 48 h after TMP-UVA (2 µM TMP, 2 h followed by 5 s irradiation) following MCM3 downregulation. Nuclear DNA is counterstained with DAPI. Scale bar, 25 µm. **F**. Dot plot shows the distribution and mean (horizontal red line) of FANCD2 foci number per cell in each condition. Pool of three replicates are represented (n ≥100 cells per condition). Statistical analysis was conducted with Kruskal-Wallis test and Dunns post-test. *, p<0.05; ***, p<0.001. Histograms (bottom panel) show the average foci number and SD of three assays in each case. Circle dots in each column represent the values of individual replicates. Statistical analysis was conducted with one-way ANOVA followed by Bonferroni post-test. **, p<0.01; ns, not significant.

MCM3 downregulation also enhanced the phenotype of persistent FANCD2 foci 48 h after TMP-UVA in PRIMPOL KO cells (**Figure 6E-F**), reflecting their dependency on backup origins to establish new replication forks and maximize fork convergence at ICLs.

## DISCUSSION

Re-priming of DNA synthesis downstream of lesions in the template is now recognized as a prominent mechanism of DNA damage tolerance (DDT), next to those involving translesive synthesis or the temporary invasion of the complementary strand (template switch). DDT pathways mediate the passage of replication forks through damaged DNA, avoiding interruptions in DNA synthesis that could increase genome instability and leaving the lesions to be repaired post-replicatively (reviewed by Branzei and Szakal, 2016; Muñoz and Méndez, 2017; Vaisman and Woodgate, 2017; Pilzecker et al., 2019).

ICLs are amongst the most cytotoxic DNA lesions and, at least in theory, represent an absolute roadblock for both replication and transcription. This fits with the model in which ICL repair is initiated by the convergence of two forks arriving from opposite sides at the lesion (Zhang et al., 2015a). However, other studies have shown that at least in avian and mammalian cells, the first fork arriving at the lesion may traverse the ICL (Huang et al., 2013, 2019; Ling et al., 2016; Mutreja et al., 2018). Both scenarios (fork convergence and traverse) result in similar X-shaped replicated DNA structures that can be processed for repair by the FA pathway. The use of one or the other may depend on the distance between adjacent origins, which is proportional to the time needed for a second fork to reach the ICL after the first one has been stalled, or by the availability of a primase to reinitiate DNA synthesis downstream of the lesion (see model in **Figure 7**).

**Figure 7.**
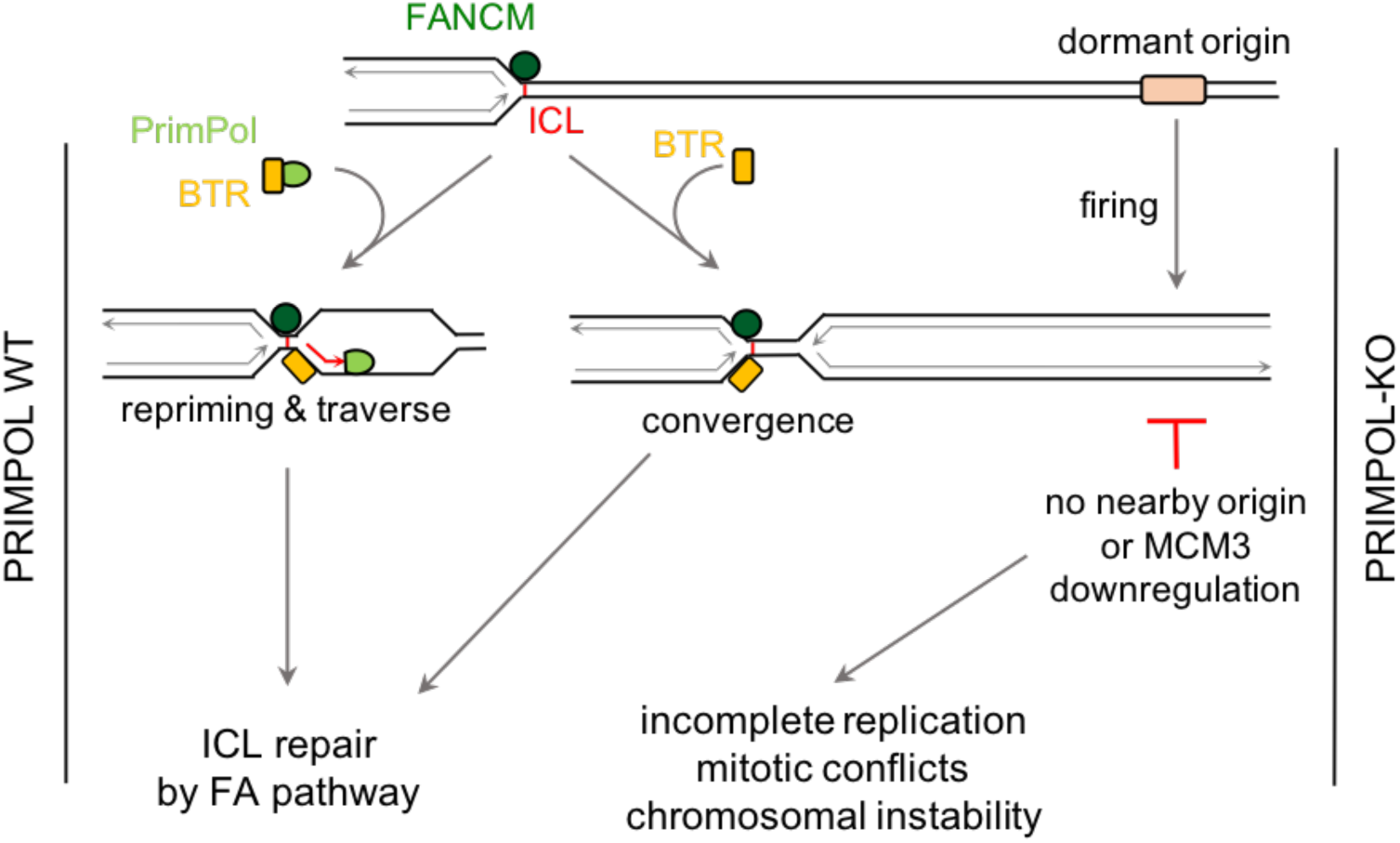
Role of PrimPol in ICL traverse and its consequences for ICL repair: a model. In PRIMPOL-proficient cells, BTR may promote its binding to ICLs. The synthesis of a new primer molecule downstream of the lesion likely disrupts the PrimPol-BTR interaction. PrimPol promotes continuous DNA synthesis and iniciation of ICL repair. In the absence of PrimPol, ICL traverse is much less efficient and cells rely on dormant origin activation to promote fork convergence at the ICL. In origin-poor regions or when dormant origin availability is limited by MCM downregulation, a fraction of the ICLs remain unrepaired, generating chromosomal instability and eventual cell death. See text for further details.

The molecular mechanisms behind replisome translocation in the ICL traverse are starting to be elucidated. FANCM translocase interacts with BTR (BLM/TOP3A/RMI1-2; Deans and West, 2009; Hoadley et al., 2012) and the traverse reaction becomes less efficient if BLM is inactive or the BTR-FANCM interaction is disrupted (Ling et al., 2016). FANCM also interacts with replisome components, including PCNA (Rohleder et al., 2016) and MCM, the core component of the CMG helicase (Huang et al., 2019). The FANCM-MCM interaction is mediated by FANCD2 and requires ATR function. Proximity ligation microscopy assays have shown that ATR and FANCD2 trigger the release of the GINS component of CMG, presumably to loose the MCM hexameric ring and facilitate its passage through the ICL (Huang et al., 2019). Of note, the ability of helicases to operate across an ICL has an interesting antecedent in bacterial DNAB helicase/translocase (Bastia et al., 2008).

Coupled to the relocation of the replisome, a primer needs to be synthesized downstream of the ICL to provide a new starting point in the leading strand for replicative polymerase Polε. We have identified PrimPol as the main primase involved in this step. PrimPol is redistributed to chromatin upon exposure to DNA crosslinking agents MMC and TMP-UVA. The recruitment of PrimPol to the vicinity of ICLs is likely mediated by RPA, which performs this role at other polymerase-stalling lesions (Wan et al., 2013; Guilliam et al., 2015, 2017). RPA could be assisted by BTR, which recognizes ICLs and interacts with both RPA (Wu et al., 2018) and PrimPol (this study). The fact that the PrimPol-BTR interaction is reduced during the response to MMC has also been reported for the interaction between RPA and BLM (Wu et al., 2018) and it may reflect that PrimPol is rapidly released from its interacting factors to synthesize the new primer downstream of the ICL (**Figure 7**).

The genetic ablation of PRIMPOL caused a striking reduction in the frequency of traverse reactions, but did not prevent them completely. Because none of the known TLS polymerases can synthesize DNA through ICLs (reviewed by Roy and Schärer, 2016), we favor the simple explanation that in the absence of PrimPol, the replicative Polα/Primase complex partially covers the DDT function. It is noteworthy that biochemical reconstitution of a yeast replication fork stalled at an UV-photoproduct shows that re-priming mediated by Polα/Primase is possible but inefficient in the leading strand (Taylor and Yeeles, 2018). Fungi do not have PrimPol, but most other eukaryotes have kept a second primase (PrimPol) through evolution, probably as a more efficient alternative for the re-priming reactions in DNA damage tolerance mechanisms.

In PRIMPOL KO cells, the impairment of traverse reactions was partially compensated by a higher frequency of fork convergence events. Because PrimPol mediates fork progression across endogenous DNA obstacles (Schiavone et al., 2016; Šviković et al., 2018), its downregulation triggers the activation of backup origins (Mourón et al., 2013; Rodriguez-Acebes et al., 2018). In the context of ICL repair, the establishment of new forks from these origins increases the chance of two forks converging at the crosslink. Consistent with this notion, the downregulation of MCM proteins to prevent dormant origin activation further sensitized PRIMPOL KO cells to ICL-inducing agents (see model in **Figure 7**).

The possibility of targeting DDT proteins as an adjuvant to standard chemotherapy is currently under investigation. For instance, the combination of cisplatin with a chemical inhibitor of Polζ markedly suppressed tumor growth of human melanoma xenografts in mice (Wojtaszek et al., 2019). Our results hint at the possibility of targeting PrimPol to enhance the efficacy of crosslinking-based chemotherapy. As shown in a recent study, PrimPol expression is upregulated in response to multiple doses of cisplatin as part of an adaptive response in BRCA1-deficient cells that have already lost the ability to stabilize stalled forks (Quinet et al., 2020). Hence, targeting PrimPol to prevent the tolerance of chemotherapy-induced ICLs could be clinically relevant in BRCA1/2-deficient tumors being treated with cisplatin or MMC. Future studies will determine the effectiveness of these approaches, which could be complemented by preventing the activation of backup origins using Cdc7 kinase inhibitors (Iwai et al., 2019; Rainey et al., 2017).

## METHODS

### Cell lines and manipulations

U2OS-shPRIMPOL cell line has been described (Mourón et al., 2013). Cells were grown in DMEM supplemented with 10% FBS plus penicillin-streptomycin and tested monthly for mycoplasma contamination. In U2OS-shPRIMPOL cells, expression of the shRNA targeting PRIMPOL was induced with 1 μg/ml doxycycline (Dox). U2OS PRIMPOL KO cells were generated as described (Ann et al., 2013) with minor modifications. U2OS cells were transfected with pSpCas9(BB)-2A-GFP plasmid (Addgene #48138) containing a short guide RNA targeting exon 7 in PRIMPOL gene, built with sgPRIMPOL_Fw (5’-CACCGGAGGTGCTTCTGAAAAATG) and sgPRIMPOL_Rv (5’-AAACCATTTTTCAGAAGCACCTCC) oligonucleotides. After 24h, transfected cells were seeded at low confluency to allow single-clone colony formation. Isolated colonies were tested for PrimPol expression. To test putative PRIMPOL KO clones, the CRISPR/Cas9 target site was amplified with specific primers PRIMPOL_seq_Fw (5’-GACCTTAAGATGCGGTGTGT) and PRIMPOL_seq_Rv (5’-TGAACTCACTGGTTGCATCG), cloned into pJET1.2/blunt vector (ThermoFisher, K1232) and sequenced. To reintroduce PrimPol protein in WT or KO cells, pcDNA3.1/nV5 plasmids encoding WT, AxA, ΔZn and CH PRIMPOL derivatives (Mourón et al., 2013) were transiently transfected using Lipofectamine 2000 (Invitrogen). To induce MCM3 downregulation, cells were transfected twice with 100 nM siRNA targeting MCM3 (Ibarra et al., 2008). Mitomycin C was obtained from Panreac (#A2190); trimethylpsoralen (TMP) was from Sigma (#T6137). UVA irradiation was performed in a BS-02 irradiation chamber (Opsytec Dr. Gröbel, Ettlingen, Germany). Cell viability assays were conducted in 96-well plates. Upon treatment with the indicated drug for the indicated amount of time, viability was measured with CellTiter-Glo luminescent assay (Promega).

### Flow cytometry analyses of DNA content and BrdU incorporation

10 μM 5-bromo 2’-deoxyuridine (BrdU; Sigma #B9285) was added to the cell medium for 30 min before harvesting. Cells were fixed with ice-cold 70% ethanol for 2 h and incubated with 2M HCl for 20 min. BrdU was immunolabeled with FITC BrdU Flow Kit (BD Bioscience #559619). DNA was stained with 50 µg/ml propidium iodide (PI; Sigma # P4864). Cells were processed in a FACSCanto cytometer (BD Biosciences) and data was analyzed with FlowJo software (Tree Star Inc).

### Confocal fluorescence microscopy

For 5-ethynyl 2’-deoxyuridine (EdU) detection, cells were grown on μCLEAR polylysine-treated plates (Greiner Bio-One) and pulse-labeled with 10 µM EdU for 30 min. Labeled cells were fixed with 4% paraformaldehyde (PFA) for 10 min and permeabilized with 0.2% Triton X-100 in PBS for 5 min on ice. EdU incorporation was visualized with Click-it EdU Alexa Fluor 647 flow cytometry assay (ThermoFisher). Nuclear DNA was stained with 100 µg/ml 4’,6-Diamidino-2-phenylindole (DAPI; Sigma #D9542). Images were acquired in an Opera High-Content Screening System with an APO 20x, 0.7 numerical aperture (NA) objective, and analyzed with Acapella software (PerkinElmer).

For FANCD2 detection, cells growing on μCLEAR polylysine-treated plates were treated before fixation with 0.5% Triton X-100 in CSK buffer (10 mM Pipes-KOH pH 7.0, 100 mM NaCl, 300 mM sucrose, 3 mM MgCl_2_) to pre-extract soluble proteins. Cells were fixed in 4% PFA for 10 min and incubated with blocking solution (3% bovine serum albumin (BSA), 0.05% Tween-20 in PBS) for 30 min. Incubations with anti-FANCD2 primary antibody and the corresponding secondary antibody (**Table 1**) were performed sequentially for 1h at RT. Cell nuclei were counterstained with DAPI as before. Images were acquired in a Leica-TCS SP5-MP confocal microscope with a HCX PL APO 40x 1.4 NA oil-immersion objective and LAS AF (v. 2.5.1) software. Images were analyzed with Definiens software (PerkinElmer).

**Table 1.** Antibodies used in this study.

| **Antibodies** | **Source** | **Reference** | **Application**<br>**(Dilution)** |
| --- | --- | --- | --- |
| anti-PrimPol | Generated<br>in house | Mourón et al., 2013 | WB (1:1,000)<br>IP |
| anti-PrimPol | Generated<br>in house | This work | WB (Serum) |
| anti-CTFC | Millipore | 07-729 | WB (1:1,000) |
| anti-TUBULIN | Sigma | T6199 | WB (1:1,000) |
| anti-MEK2 | BD<br>Biosciences | 610236 | WB (1:1,000) |
| anti-FANCD2 | Fanconi<br>Anemia<br>Research<br>Found<br>(FARF) | B9699 | IF (1:100) |
| anti-FANCD2 | Novus<br>Biologicals | NB100-182 | WB (1:10,000) |
| anti-BLM | Bethyl<br>Laboratories | A300-110A | WB (1:1,000) |
| anti-RMI1 | Abcam | Ab70525 | WB (1:1,000) |
| anti-RMI2 | Novus<br>Biologicals | NBP1-89962 | WB (1:1,000) |
| anti-TOP3A | Proteintech | 14525-1-AP | WB (1:1,000) |
| anti-RPA2 | Cell<br>Signaling | 2208S | WB (1:1,000) |
| anti-MCM3 | Méndez Lab | Ibarra et al., 2008 | WB (1:1,000) |
| anti-V5 | Invitrogen | 46-0705 | WB (1:1,000) |
| anti-SMC1 | Losada's lab | Kojic et al., 2018 | WB (1 ug/mL) |
| anti-Ki67 | Cell<br>Signaling | 12202 | IHC |
| anti-CldU (rat monoclonal anti-BrdU) | Abcam | ab6326 | IF (1:100) |
| anti-IdU (mouse monoclonal anti-BrdU) | BD biosciences | 347580 | IF (1:100) |
| anti-ssDNA | Millipore | MAB3034 | IF (1:300) |
| anti-Dig | Abcam | ab76907 | IF (1:1000) |
| anti-rabbit IgG AF-488 (chicken) | Invitrogen Molecular Probes | A-21441 | IF (1:300) |
| anti-rat IgG AF-594 (goat) | Invitrogen Molecular Probes | A-11007 | IF (1:300) |
| anti-mouse IgG2a AF-647 (goat) | Invitrogen Molecular Probes | A-21241 | IF (1:300) |
| anti-goat IgG AF-647 (donkey) | Invitrogen Molecular Probes | A-21447 | IF (1:1500) |
| Horseradish peroxidase–linked ECL anti-rabbit IgG | GE Healthcare | NA934 | WB (1:5,000) |
| Horseradish peroxidase–linked ECL anti-mouse IgG | GE Healthcare | NA931 | WB (1:5,000) |
| Horseradish peroxidase–linked ECL anti-rat IgG | GE Healthcare | NA935 | WB (1:5,000) |

### Biochemical fractionation, immunoprecipitation and immunoblots

Whole cell extracts were prepared by suspension of cells in Laemmli buffer (50mM Tris-HCl pH 6.8, 10% glycerol, 3% SDS, 0.006 w/v bromophenol blue and 5% 2-mercaptoethanol) followed by 30 sec sonication in a Branson Digital Sonifier at 15% amplitude. Biochemical fractionation was performed as described (Mendez and Stillman, 2000). SDS-PAGE, protein transfer to nitrocellulose and immunoblots were performed using standard protocols. For immunoprecipitation assays, either 5 × 10^6^ (for IP-immunoblot) or 20 × 10^6^ cells (for mass spectrometry) were lysed in NP40 buffer (150 mM NaCl, 50 mM Tris pH8, 1% NP40, 0.2 mM PMSF, 5 mM NaF, 5 mM NaVO4, 1 mM DTT) supplemented with protease inhibitors (Roche # 11836153001). Anti-PrimPol antibody (Table 1) was coupled to magnetic M-270 Epoxy Dynabeads® (7.5 μg antibody/ mg beads; Life Technologies), and IPs were performed following manufacturer instructions. Cell lysates were incubated with 1.5 mg antibody-coupled beads for 45 min at 4º C. Protein-antibody-bead complexes were washed 3x with lysis buffer and collected for immunoblot or mass spectrometry assays.

### Mass spectrometry

PrimPol immunoprecipitates were eluted in 8M urea, 100 mM Tris/HCl pH 8.0. Eluates were incubated with 15 mM Tris(2-carboxyethyl)phosphine (TCEP), 45 mM chloroacetamide for 1h at 25 ºC in the dark. Protein samples digested with 100 ng of endopeptidase LysC for 4h/ 25 ºC and subsequently with 100 ng of trypsin (16h/ 37 ºC). Samples were analyzed in LTQ Orbitrap Velos (Thermo Fisher Scientific) coupled to a nanoLC Ultra 1D+ system (Eksigent), 5 µL of each sample were loaded onto a reversed-phase C18, 5 µm, 0.1 × 20 mm trapping column (NanoSeparations) and washed for 10 min at 2.5 µl/min with 0.1 % FA. The peptides were eluted at a flow rate of 250 nl/min onto an analytical column packed with ReproSil-Pur C18-AQ beads, 2.4 μm, 75 μm x 50 cm, heated to 45°C. Solvent A was 4% ACN in 0.1% FA and Solvent B acetonitrile in 0.1% FA. Peptides were separated using a lineal gradient from 4% to 33% B in 100 min. Total run time, including washing and re-equilibrating, was 120 min.

Data was analyzed with MaxQuant (Cox and Mann, 2008) v1.5.3.30 with Andromeda (Cox et al., 2011) as the search engine against a *Homo sapiens* database supplemented with the sequences of the contaminants most frequently detected in the proteomics lab (UniProtKB/Swiss-Prot, 20,599 sequences). Minimal peptide length was set to 6 amino acids and a maximum of two missed cleavages were allowed. Peptides and proteins were filtered at 1% false discovery rate (FDR). Student’s T-test statistics were used for determining the significance of the intensity of each protein.

### Single-molecule analysis of DNA replication in stretched fibers

To measure fork progression rate and frequency of origin activation, cells were pulse-labeled sequentially with 50 μM 5-Chloro-2’-deoxyuridine (CldU; 20 min) followed by 250 μM 5-iodo-2’-deoxyuridine (IdU; 20 min). Labeled cells were harvested and resuspended in 0.2 M Tris pH 7.4, 50 mM EDTA, 0.5% SDS. To evaluate fork restart, cells were pulsed with CldU for 20 min, treated with 50 µg/mL MMC for 30 min and pulsed with IdU for 20 min in fresh medium. In all cases, stretched DNA fibers were prepared as described (Mourón et al., 2013). For immunodetection of labeled tracks, fibers were incubated with anti-CldU and anti-IdU primary antibodies (1h/ RT) and the corresponding secondary antibodies (30 min/RT) in a humidity chamber. Fiber integrity was assessed by staining with anti-ssDNA. Microscopy images were obtained in a DM6000 B Leica unit equipped with an HCX PL APO 40x, 0.75 NA objective. A standard conversion rate of 1 μm = 2.59 kb was used (Jackson and Pombo, 1998). For the visualization of ICLs in DNA fibers we used a recent variation (Mutreja et al., 2018) of the original protocol (Huang et al., 2013). In brief, cells were incubated with 5 μM Dig-TMP in phenol-free, FBS-free, Pen-Strep-free DMEM media for 1h in the dark and irradiated with UV-A (3J/cm^2^). Fibers were incubated anti-Dig antibody for 2h and the corresponding secondary antibody for 1h.

### Chromosome stability

Cells in culture were treated with 0.1 μg/ml colcemide for 4 h, harvested by trypsinization, incubated in 75 mM KCl (30 min/ 37°C), fixed in 3:1 methanol:acetic acid and dropped onto slides to obtain chromosome spreads. Chromosomes were stained with Leishman stain (Sigma #L6254) and mounted with ProLong-Gold (Invitrogen).

### Sensitivity of mouse strains to MMC and histopathology analyses

Mice were housed at the CNIO Animal Facility in accordance with “Federation for Laboratory Animal Science Associations” (FELASA) guidelines. All animal procedures were approved by the Institutional Animal Care and Use Committee (IACUC) from *Instituto de Salud Carlos III* (Spain). PRIMPOL KO mice have been described (García-Gómez et al., 2013; Mourón et al., 2013). For survival assays, cohorts of 9-15 mice of both sexes (8 to 14 weeks-old) were injected with MMC at either 7.5, 10 or 15 mg/kg of body weight. Survival was monitored for two weeks. Mice were sacrificed if they reached the humane endpoint as defined in the ethical permit. For histopathology analyses, age- and gender-matched pairs of mice from WT or KO genotype were injected with 10 mg/kg MMC. Mice tissues were fixed in 10% buffered formalin (Sigma) and embedded in paraffin using standard procedures. 3 µm sections were stained with hematoxylin and eosin (H&E). Ki-67 staining was performed in an automatic Ventana Discovery XT platform (Roche). Tissue slides were digitalized in a Mirax scan or Axio Scan.Z1 (Carl Zeiss). Staining was analyzed using AxioVision digital image software (Carl Zeiss). Areas of positive staining were normalized to the total cellular area in the tissue.

## Acknowledgements

We thank all members of the CNIO DNA Replication and Chromosome Dynamics Groups as well as Kurt Jacobs and Jana Krietsch (Lopes lab) for discussions and useful comments on the manuscript. We are grateful to Diego Megías and Sandra Rodríguez, Heads of the CNIO Confocal Microscopy and Cytogenetics Units, respectively, for their excellent technical assistance. The Fanconi Anemia Research Fund (FARF, http://www.fanconi.org) kindly provided anti-FANCD2 antibody. Research was supported by the Spanish Ministry of Science, Innovation and Universities (grants BFU2013-49153P and BFU2016-80402R to JM and PGC2018.093576-B-C21 to LB, co-sponsored by ERDF funds from the EU), the Swiss National Science Foundation (grant 31003A_169959 to ML) and an ERC Consolidator Grant (617102 to ML).

## Author contributions

DG-A performed most experiments, with help from EB-R (confocal microscopy), SL (flow cytometry), KM (single-molecule analyses of ICL traverse), and SM (PrimPol immunoprecipitation). FG and J Muñoz carried out mass spectrometry analyses. JM designed and supervised the study, with contributions from LB and ML. DG-A and JM wrote the manuscript.

## REFERENCES

Akkari, Y.M.N., Bateman, R.L., Reifsteck, C.A., Olson, S.B., and Grompe, M. (2000). DNA Replication Is Required To Elicit Cellular Responses to Psoralen-Induced DNA Interstrand Cross-Links. Mol. Cell. Biol. 20, 8283–8289.

Ann, F R., Hsu, P.D., Wright, J., Agarwala, V., Scott, D.A., and Zhang, F. (2013). Genome engineering using the CRISPR-Cas9. Nat. Protoc. 8, 2281–2308.

Bastia, D., Zzaman, S., Krings, G., Saxena, M., Peng, X., and Greenberg, M.M. (2008). Replication termination mechanism as revealed by Tus-mediated polar arrest of a sliding helicase. Proc. Natl. Acad. Sci. U. S. A. 105, 12831–12836.

Bianchi, J., Rudd, S.G., Jozwiakowski, S.K., Bailey, L.J., Soura, V., Taylor, E., Stevanovic, I., Green, A.J., Stracker, T.H., Lindsay, H.D., et al. (2013). Primpol Bypasses UV Photoprodcts during Eukaryotic Chomosomal DNA Replication. Mol. Cell 52, 566–573.

Branzei, D., and Szakal, B. (2016). DNA damage tolerance by recombination: Molecular pathways and DNA structures. DNA Repair (Amst) 44, 68–75.

Ceccaldi, R., Sarangi, P., and D’Andrea, A.D. (2016). The Fanconi anaemia pathway: New players and new functions. Nat. Rev. Mol. Cell Biol. 17, 337–349.

Ciccia, A., Ling, C., Coulthard, R., Yan, Z., Xue, Y., Meetei, A.R., Laghmani, E.H., Joenje, H., McDonald, N., de Winter, J.P., et al. (2007). Identification of FAAP24, a Fanconi Anemia Core Complex Protein that Interacts with FANCM. Mol. Cell 25, 331–343.

Cox, J. and Mann, M. (2008). MaxQuant enables high peptide identification rates, individualized p.p.b.-range mass accuracies and proteome-wide protein quantification. Nat. Biotechnol. 26, 1367–1372.

Cox, J., Neuhauser, N., Michalski, A., Scheltema, R.A., Olsen, J. V., and Mann, M. (2011). Andromeda: A peptide search engine integrated into the MaxQuant environment. J. Proteome Res. 10, 1794–1805.

Deans, A.J., and West, S.C. (2009). FANCM Connects the Genome Instability Disorders Bloom’s Syndrome and Fanconi Anemia. Mol. Cell 36, 943–953.

Deans, A.J., and West, S.C. (2011). DNA interstrand crosslink repair and cancer. Nat. Rev. Cancer 11, 467–480.

Dendouga, N., Gao, H., Moechars, D., Janicot, M., Vialard, J., and McGowan, C.H. (2005). Disruption of Murine Mus81 Increases Genomic Instability and DNA Damage Sensitivity but Does Not Promote Tumorigenesis. Mol. Cell. Biol. 25, 7569–7579.

Dronkert, M.L.G., de Wit, J., Boeve, M., Vasconcelos, M.L., van Steeg, H., Tan, T.L.R., Hoeijmakers, J.H.J., and Kanaar, R. (2000). Disruption of Mouse SNM1 Causes Increased Sensitivity to the DNA Interstrand Cross-Linking Agent Mitomycin C. Mol. Cell. Biol. 20, 4553–4561.

García-Gómez, S., Reyes, A., Martínez-Jiménez, M.I., Chocrón, E.S., Mourón, S., Terrados, G., Powell, C., Salido, E., Méndez, J., Holt, I.J., Blanco, L. (2013). PrimPol, an Archaic Primase/Polymerase Operating in Human Cells. Mol. Cell 52, 541–553.

Garcia-Higuera, I., Taniguchi, T., Ganesan, S., Meyn, M.S., Timmers, C., Hejna, J., Grompe, M., and D’Andrea, A.D. (2001). Interaction of the Fanconi anemia proteins and BRCA1 in a common pathway. Mol. Cell 7, 249–262.

Ge, X.Q., Jackson, D.A., and Blow, J.J. (2007). Dormant origins licensed by excess Mcm2-7 are required for human cells to survive replicative stress. Genes Dev. 21, 3331–3341.

Guilliam, T.A., Jozwiakowski, S.K., Ehlinger, A., Barnes, R.P., Rudd, S.G., Bailey, L.J., Skehel, J.M., Eckert, K.A., Chazin, W.J., and Doherty, A.J. (2015). Human PrimPol is a highly error-prone polymerase regulated by single-stranded DNA binding proteins. Nucleic Acids Res. 43, 1056–1068

Guilliam, T.A., Brissett, N.C., Ehlinger, A., Keen, B.A., Kolesar, P., Taylor, E.M., Bailey, L.J., Lindsay, H.D., Chazin, W.J., and Doherty, A.J. (2017). Molecular basis for PrimPol recruitment to replication forks by RPA. Nat. Commun. 8, 15222

Hoadley, K.A., Xue, Y., Ling, C., Takata, M., Wang, W., and Keck, J.L. (2012). Defining the molecular interface that connects the Fanconi anemia protein FANCM to the Bloom syndrome dissolvasome. Proc. Natl. Acad. Sci. U. S. A. 109, 4437–4442.

Huang, J., Liu, S., Bellani, M.A., Thazhathveetil, A., Ling, C., deWinter, J.P., Wang, Y., Wang, W., and Seidman, M.M. (2013). The DNA Translocase FANCM/MHF Promotes Replication Traverse of DNA Interstrand Crosslinks. Mol. Cell 52, 434–446.

Huang, J., Zhang, J., Bellani, M.A., Li, L., Wang, W., and Seidman, M.M. (2019). Remodeling of Interstrand Crosslink Proximal Article Remodeling of Interstrand Crosslink Proximal Replisomes Is Dependent on ATR, FANCM, and FANCD2. Cell Rep. 27, 1794–1808.

Ibarra, A., Schwob, E., and Méndez, J. (2008). Excess MCM proteins protect human cells from replicative stress by licensing backup origins of replication. Proc. Natl. Acad. Sci. U. S. A. 105, 8956–8961.

Iwai, K., Nambu, T., Dairiki, R., Ohori, M., Yu, J., Burke, K., Gotou, M., Yamamoto, Y., Ebara, S., Shibata, S., et al. (2019). Molecular mechanism and potential target indication of TAK-931, a novel CDC7-selective inhibitor. Sci. Adv. 5, eaav3660.

Jackson, D.A., and Pombo, A. (1998). Replicon clusters are stable units of chromosome structure: Evidence that nuclear organization contributes to the efficient activation and propagation of S phase in human cells. J. Cell Biol. 140, 1285–1295.

Knipscheer, P., Räschle, M., Smogorzewska, A., Enoiu, M., Ho, T.V., Schärer, O.D., Elledge, S.J., and Walter, J.C. (2009). The fanconi anemia pathway promotes replication-dependent DNA interstrand cross-link repair. Science 326, 1698–1701.

Kojic, A., Cuadrado, A., De Koninck, M., Giménez-Llorente, D., Rodríguez-Corsino, M., Gómez-López, G., Le Dily, F., Marti-Renom, M.A., and Losada, A. (2018). Distinct roles of cohesin-SA1 and cohesin-SA2 in 3D chromosome organization. Nat. Struct. Mol. Biol. 25, 496–504.

Liang, C.C., Zhan, B., Yoshikawa, Y., Haas, W., Gygi, S.P., and Cohn, M.A. (2015). UHRF1 Is a sensor for DNA interstrand crosslinks and recruits FANCD2 to initiate the Fanconi Anemia pathway. Cell Rep. 10, 1947–1957.

Ling, C., Huang, J., Yan, Z., Li, Y., Ohzeki, M., Ishiai, M., Xu, D., Takata, M., Seidman, M., and Wang, W. (2016). Bloom syndrome complex promotes FANCM recruitment to stalled replication forks and facilitates both repair and traverse of DNA interstrand crosslinks. Cell Discov. 2, 16047.

Lopez-Martinez, D., Liang, C.-C., and Cohn, M.A. (2016). Cellular response to DNA interstrand crosslinks: the Fanconi anemia pathway. Cell. Mol. Life Sci. 73, 3097–3114.

Lu, H., Guo, X., Meng, X., Liu, J., Allen, C., Wray, J., Nickoloff, J.A., Shen, Z. (2005). The BRCA2-interacting protein BCCIP functions in RAD51 and BRCA2 focus formation and homologous recombinational repair. Mol Cell Biol. 25, 1949–1957.

Manthei, K.A. and Keck, J.L. (2013). The BLM dissolvasome in DNA replication and repair. Cell Mol Life Sci 70, 4067–4084.

Martínez-Jiménez, M.I., Lahera, A., and Blanco, L. (2017). Human PrimPol activity is enhanced by RPA. Sci. Rep. 7, 783.

McPherson, J.P., Lemmers, B., Chahwan, R., Pamidi, A., Migon, E., Matysiak-Zablocki, E., Moynahan, M.E., Essers, J., Hanada, K., Poonepalli, A., et al. (2004). Involvement of mammalian Mus81 in genome integrity and tumor suppression. Science 304, 1822–1826.

Mendez, J., and Stillman, B. (2000). Chromatin Association of Human Origin Recognition Complex, Cdc6, and Minichromosome Maintenance Proteins during the Cell Cycle: Assembly of Prereplication Complexes in Late Mitosis. Mol. Cell. Biol. 20, 8602–8612.

Mourón, S., Rodriguez-Acebes, S., Martínez-Jiménez, M.I., García-Gómez, S., Chocrón, S., Blanco, L., and Méndez, J. (2013). Repriming of DNA synthesis at stalled replication forks by human PrimPol. Nat. Struct. Mol. Biol. 20, 1383–1389.

Muñoz, S., and Méndez, J. (2017). DNA replication stress: from molecular mechanisms to human disease. Chromosoma 126, 1–15.

Mutreja, K., Krietsch, J., Hess, J., Ursich, S., Berti, M., Roessler, F.K., Zellweger, R., Patra, M., Gasser, G., and Lopes, M. (2018). ATR-Mediated Global Fork Slowing and Reversal Assist Fork Traverse and Prevent Chromosomal Breakage at DNA Interstrand Cross-Links. Cell Rep. 24, 2629–2642.

Pilzecker, B., Buoninfante, O.A., and Jacobs, H. (2019). DNA damage tolerance in stem cells, ageing, mutagenesis, disease and cancer therapy. Nucleic Acids Res. 47, 7163–7181.

Quinet, A., Tirman, S., Jackson, J., Šviković, S., Lemaçon, D., Carvajal-Maldonado, D., González-Acosta, D., Vessoni, A.T., Cybulla, E., Wood, M., Tavis, S., Batista, L.F.Z, Méndez, J., Sale, J.E., Vindigni, A. (2020). PRIMPOL-Mediated Adaptive Response Suppresses Replication Fork Reversal in BRCA-Deficient Cells. Mol. Cell 77, 461–474.

Rainey, M.D., Quachthithu, H., Gaboriau, D., and Santocanale, C. (2017). DNA Replication Dynamics and Cellular Responses to ATP Competitive CDC7 Kinase Inhibitors. ACS Chem. Biol. 12, 1893–1902.

Räschle, M., Knipsheer, P., Enoiu, M., Angelov, T., Sun, J., Griffith, J.D., Ellenberger, T.E., Schärer, O.D., and Walter, J.C. (2008). Mechanism of Replication-Coupled DNA Interstrand Crosslink Repair. Cell 134, 969–980.

Rodriguez-Acebes, S., Mourón, S., and Méndez, J. (2018). Uncoupling fork speed and origin activity to identify the primary cause of replicative stress phenotypes. J. Biol. Chem. 293, 12855–12861.

Rohleder, F., Huang, J., Xue, Y., Kuper, J., Round, A., Seidman, M., Wang, W., and Kisker, C. (2016). FANCM interacts with PCNA to promote replication traverse of DNA interstrand crosslinks. Nucleic Acids Res. 44, 3219–3232.

Roy, U., and Schärer, O.D. (2016). Involvement of translesion synthesis DNA polymerases in DNA interstrand crosslink repair. DNA Repair (Amst) 44, 33–41.

Schiavone, D., Jozwiakowski, S.K., Romanello, M., Guilbaud, G., Guilliam, T.A., Bailey, L.J., Sale, J.E., and Doherty, A.J. (2016). PrimPol Is Required for Replicative Tolerance of G Quadruplexes in Vertebrate Cells. Mol. Cell 61, 161–169.

Šviković, S., Crisp, A., Tan-Wong, S.M., Guilliam, T.A., Doherty, A.J., Proudfoot, N.J., Guilbaud, G., and Sale, J.E. (2019). R-loop formation during S phase is restricted by PrimPol-mediated repriming. EMBO J. 38, pii: e99793.

Tay, L.S., Krishnan, V., Sankar, H., Chong, Y.L., Chuang, L.S.H., Tan, T.Z., Kolinjivadi, A.M., Kappei, D., Ito, Y. (2018). RUNX Poly(ADP-Ribosyl)ation and BLM Interaction Facilitate the Fanconi Anemia Pathway of DNA Repair. Cell Rep. 24, 1747–1755.

Taylor, M.R.G., and Yeeles, J.T.P. (2018). The Initial Response of a Eukaryotic Replisome to DNA Damage. Mol. Cell 70, 1067–1080.

Thongthip, S., Bellani, M., Gregg, S.Q., Sridhar, S., Conti, B.A., Chen, Y., Seidman, M.M., and Smogorzewska, A. (2016). Fan1 deficiency results in DNA interstrand cross-link repair defects, enhanced tissue karyomegaly, and organ dysfunction. Genes Dev. 30, 645–659.

Torregrosa-Muñumer, R., Forslund, J.M.E., Goffart, S., Pfeiffer, A., Stojkovic, G., Carvalho, G., Al-Furoukh, N., Blanco, L., Wanrooij, S., and Pohjoismäki, J.L.O. (2017). PrimPol is required for replication reinitiation after mtDNA damage. Proc. Natl. Acad. Sci. U. S. A. 114, 11398–11403.

Vaisman, A., and Woodgate, R. (2017). Translesion DNA polymerases in eukaryotes: what makes them tick? Crit. Rev. Biochem. Mol. Biol. 52, 274–303.

Wan, L., Lou, J., Xia, Y., Su, B., Liu, T., Cui, J., Sun, Y., Lou, H., and Huang, J. (2013). HPrimpol1/CCDC111 is a human DNA primase-polymerase required for the maintenance of genome integrity. EMBO Rep. 14, 1104–1112.

Wang, Y., Leung, J.W., Jiang, Y., Lowery, M.G., Do, H., Vasquez, K.M., Chen, J., Wang, W., and Li, L. (2013). FANCM and FAAP24 Maintain Genome Stability via Cooperative as Well as Unique Functions. Mol. Cell 49, 997–1009.

Wojtaszek, J.L., Chatterjee, N., Najeeb, J., Ramos, A., Lee, M., Bian, K., Xue, J.Y., Fenton, B.A., Park, H., Li, D., Hemann, M.T., Hong, J., Walker, G.C., Zhou, P. (2019). A Small Molecule Targeting Mutagenic Translesion Synthesis Improves Chemotherapy. Cell 178, 152–159.

Wu, W., Rokutanda, N., Takeuchi, J., Lai, Y., Maruyama, R., Togashi, Y., Nishikawa, H., Arai, N., Miyoshi, Y., Suzuki, N., Saeki, Y., Tanaka, K., Ohta, T. (2018). HERC2 facilitates BLM and WRN helicase complex interaction with RPA to suppress G-quadruplex DNA. Cancer Res. 78, 6371–6385.

Yan, Z., Delannoy, M., Ling, C., Daee, D., Osman, F., Muniandy, P.A., Shen, X., Oostra, A.B., Du, H., Steltenpool, J., Lin, T., Schuster, B., Décaillet, C., Stasiak, A., Stasiak, A.Z., Stone, S., Hoatlin, M.E., Schindler, D., Woodcock, C.L., Joenje, H., Sen, R., de Winter, J.P., Li, L., Seidman, M.M., Whitby, M.C., Myung, K., Constantinou, A., Wang, W. (2010). A Histone-Fold Complex and FANCM Form a Conserved DNA-Remodeling Complex to Maintain Genome Stability. Mol. Cell 37, 865–788.

Zhang, J., and Walter, J.C. (2014). Mechanism and regulation of incisions during DNA interstrand cross-link repair. DNA Repair (Amst) 19, 135–142.

Zhang, J., Dewar, J.M., Budzowska, M., Motnenko, A., Cohn, M.A., and Walter, J.C. (2015a). DNA interstrand cross-link repair requires replication-fork convergence. Nat. Struct. Mol. Biol. 22, 242–247.

Zhang, T., Wilson, A.F., Mahmood Ali, A., Namekawa, S.H., Andreassen, P.R., Ruhikanta Meetei, A., and Pang, Q. (2015b). Loss of Faap20 causes hematopoietic stem and progenitor cell depletion in mice under genotoxic stress. Stem Cells 33, 2320–2330.

