## Supplemental information for "PrimPol primase mediates replication traverse of DNA interstrand crosslinks"

### Supplementary Figure Legends

**Supplementary Figure 1** (related to Figure 1). **Sequence of PRIMPOL mutations in the alleles of U2OS PrimPol KO cells and cell synchronization for the proteomics experiments. A.** Genomic sequences derived from WT and KO alleles in U2OS-PRIMPOL KO cells obtained with CRISPR/Cas9. The Cas9 guide RNA sequence is marked in red. As the 4q35.1 chromosome locus is amplified in U2OS cells, three PRIMPOL alleles were targeted to achieve a full KO. **B.** Flow cytometry profiles of DNA content (PI) in the synchronized cell cultures used for PrimPol IP and mass spectrometry. The percentage of cells in each phase of the cell cycle is indicated. **C.** List of enrichment ratios and statistical significance (T-test -log p-value) of selected proteins identified in mass spectrometry in the different conditions. In enrichment columns, green and red colored numbers represent statistically significant and not-significant comparisons, respectively.

**Supplementary Figure 2** (related to Figure 2). **PrimPol downregulation or ablation impairs cell proliferation and DNA synthesis in the presence of ICLs. A.** CellTiter-Glo viability assays in U2OS-shPRIMPOL cells 6 days after treatment with 1 µg/ml MMC (2 h) or TMP-UVA (2 µM TMP, 2 h followed by 5 s irradiation). When indicated, PRIMPOL expression was downregulated by a specific shRNA. Histograms represent the total number of viable cells (average and SD of three assays) at the end of the experiment, expressed as percentage of the number of untreated cells. Circle dots in each column represent the values of individual replicates. Statistical analysis was conducted with one-way ANOVA and Bonferroni post-test. *, $p<0.05$; ***, $p<0.001$. Immunoblots (bottom) show the levels of PrimPol in control cells or after the expression of shPRIMPOL for 3 or 6 days. Ponceau-S staining is shown as loading control. **B.** Flow cytometry profiles of BrdU incorporation (y-axis) vs DNA content (PI; x-axis) in WT (blue) and PRIMPOL KO (orange) cells, either collected without treatment (UNT) or at the indicated times after exposure to UVA (5 s irradiation). The DNA content profiles are shown on top of the boxes (cell count). Dashed horizontal lines inside panels are included for comparative purposes between WT and KO BrdU incorporation levels. **C.** Representative confocal microscopy images of EdU staining in WT and PRIMPOL KO cells, 8 h after UVA (control) or TMP-UVA treatment (2 µM TMP, 2h followed by 5 s irradiation). Nuclear DNA is counterstained with DAPI. When indicated, V5-PrimPol was reintroduced into KO cells. Scale bar, 25 µm. **D.** Dot plots indicate the distribution of nuclear EdU intensity, and the mean values are indicated by horizontal red lines. Data from the combination of three replicates (n ≥200 cells/ condition). Statistical analysis was conducted with Kruskal-

Wallis test and Dunns post-test. ***, $p<0.001$. Histograms (right panel) represent the median EdU intensity for each condition, normalized to UVA-treated controls (average and SD of three replicates). Circle dots in each column represent the values of individual replicates. Statistical analysis was conducted with one-way ANOVA and Bonferroni post-test. *, $p<0.05$; ns, not significant. **E.** Immunoblots show levels of endogenous PrimPol and exogenous (V5) PrimPol expressed in PRIMPOL KO cells. Tubulin is shown as loading control.

**Supplementary Fig 3** (related to Figure 3). **PrimPol promotes fork restart upon MMC treatment. A.** Representative images of DNA fibers derived from WT and KO cells. The positions of restarted forks are indicated with yellow arrows. **B.** Immunoblots showing expression levels of V5-PrimPol proteins (WT and mutant derivatives). Ponceau-S staining is shown as loading control.

**Supplementary Figure 4** (related to Figure 4). **Delayed clearance of FANCD2 foci in PRIMPOL KO cells. A.** Confocal IF microscopy images of FANCD2 staining at the indicated times in control or MMC-treated (0.5 µg/ml; 2 h) WT and PRIMPOL KO cells. Nuclear DNA is counterstained with DAPI. Scale bar, 25 µm. **B.** Dot plot shows the distribution and mean (horizontal red line) of FANCD2 foci number per cell in each condition. Pool of three replicates are represented ($n \geq 100$ cells per condition). Statistical analysis was conducted with Kruskal-Wallis test and Dunns post-test. ***, $p<0.001$. Histograms (bottom panel) show the average foci number and SD of three assays in each case. Circle dots in each column represent the values of individual replicates. Statistical analysis was conducted with one-way ANOVA followed by Bonferroni post-test. *, $p<0.05$; **, $p<0.01$.

**A**

4q35.1 - PRIMPOL exon 7

sgRNA target

WT CCATGGATTTCCCCATTTTTCAGAAGCACCTGCAAG

KO allele 1 CCA----------------------------------------GAAGCACCTGCAAG
KO allele 2 CCATGGATTTCCCCATTTT---CAGAAGCACCTGCAAG
KO allele 3 CCATGGATTTCCCCAT---CTTCAGAAGCACCTGCAAG

**B**

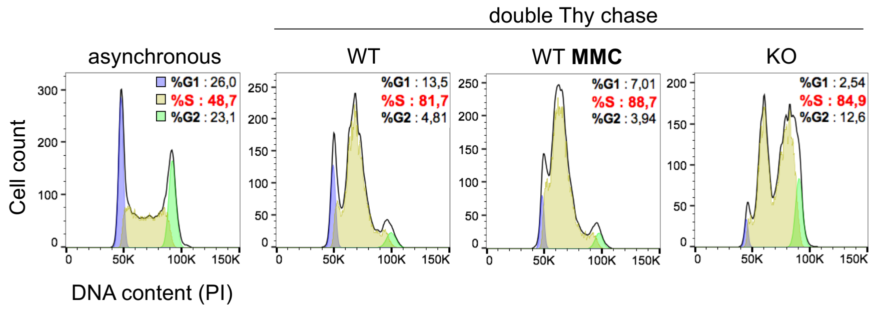


**C**

| Gene names | Enrichment WT / KO (Log2) | p-value WT vs KO (-Log) | Enrichment MMC / WT (Log2) | p-value MMC vs WT (-Log) |
|---|---|---|---|---|
| PrimPol | 6,76 | 2,44 | 0,03 | 0,12 |
| RPA1 | 1,82 | 3,45 | -0,66 | 2,40 |
| RPA3 | 2,26 | 3,61 | -0,77 | 0,82 |
| RPA2 | 2,19 | 3,20 | -0,55 | 1,82 |
| BLM | 1,74 | 2,78 | -1,38 | 2,43 |
| RMI1 | 2,68 | 3,40 | -2,23 | 3,05 |
| RMI2 | 2,05 | 2,41 | -1,74 | 2,41 |
| MHF1 | 3,99 | 1,88 | -5,16 | 4,30 |

Gonzalez-Acosta Supp Figure 2

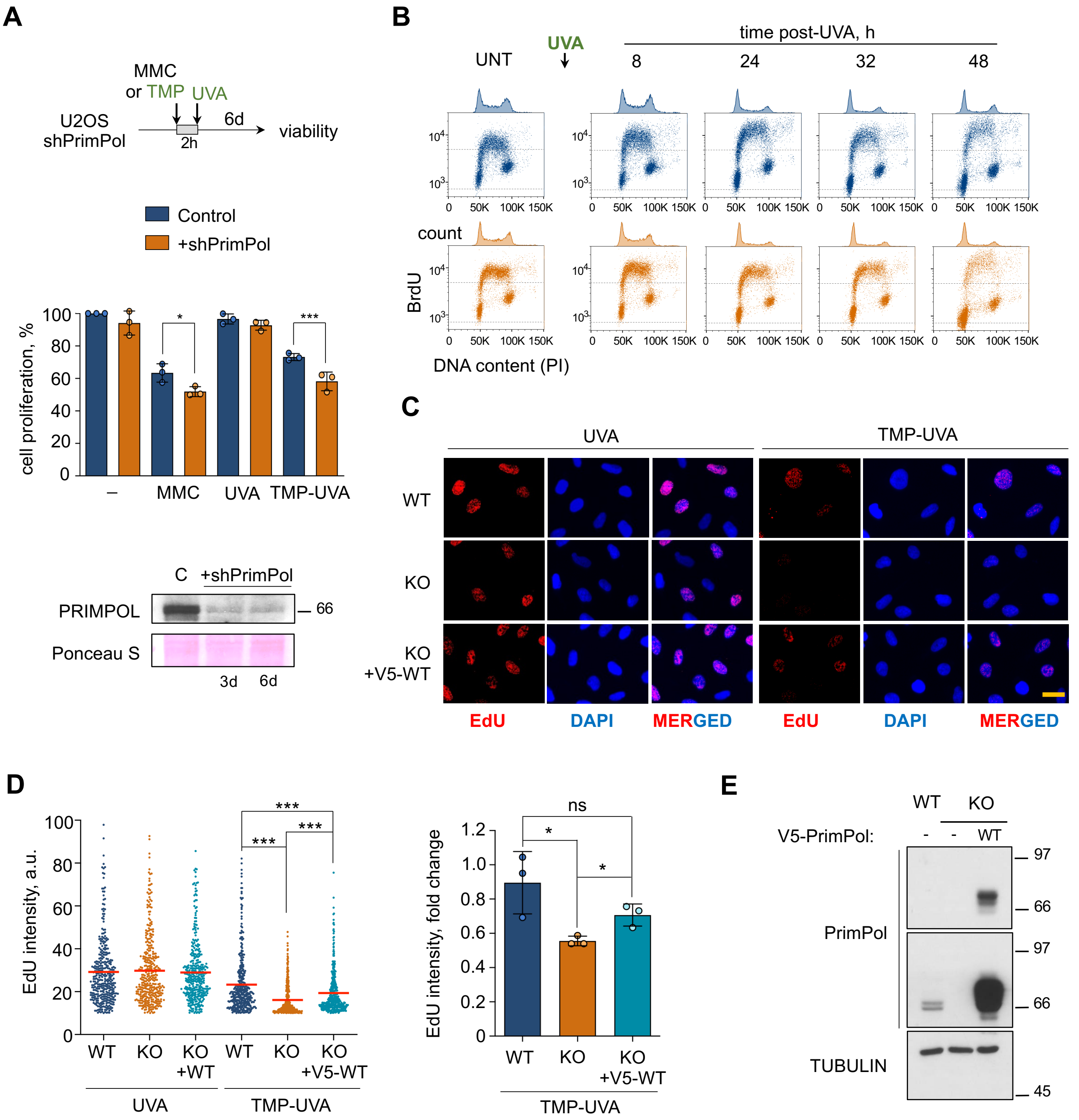

Gonzalez-Acosta Supp Figure 3

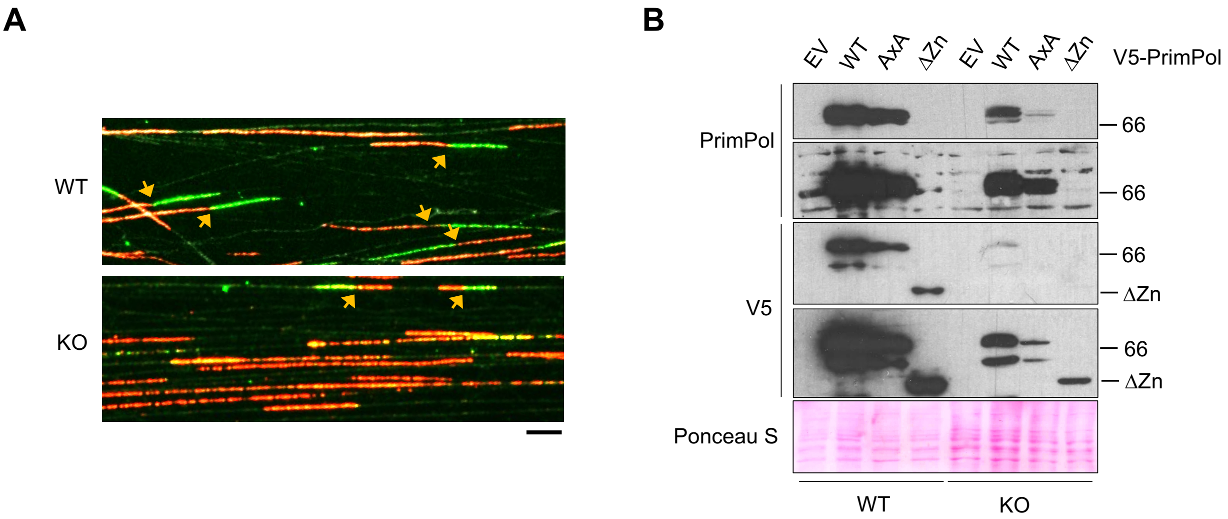

Gonzalez-Acosta Supp Figure 4

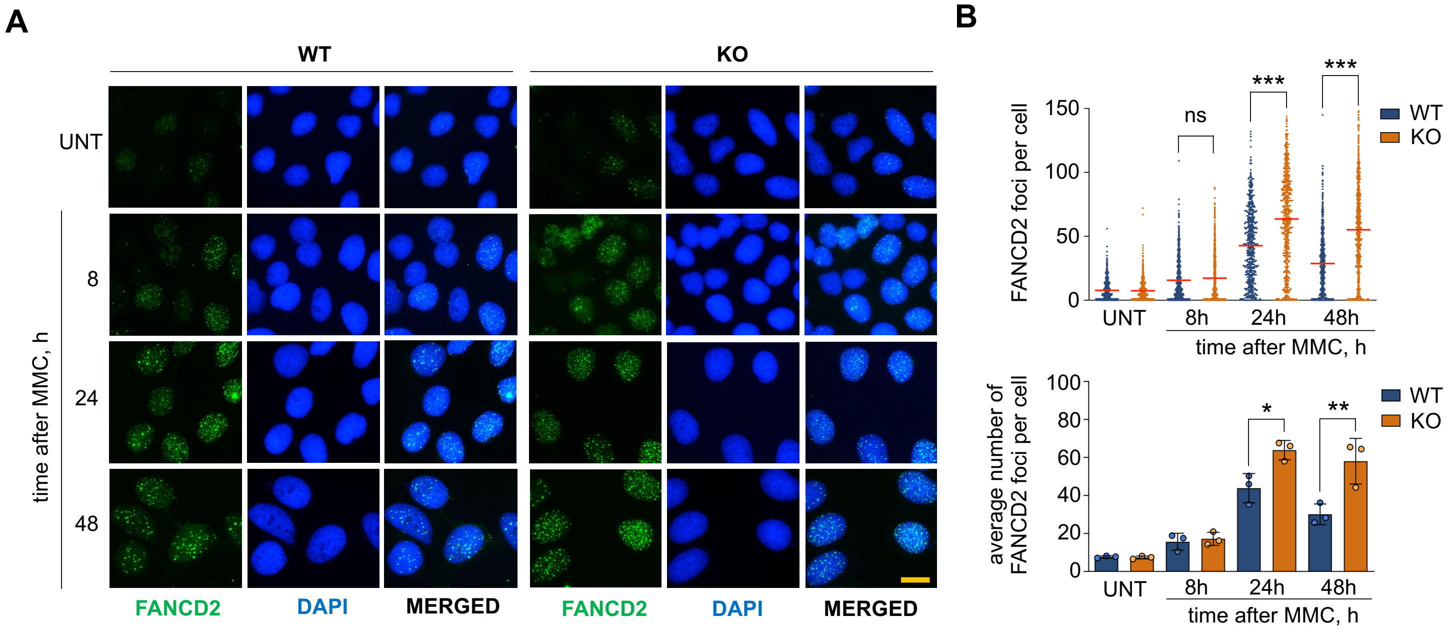
